# Exploring the impact of terminators on transgene expression in *Chlamydomonas reinhardtii* with a synthetic biology approach

**DOI:** 10.1101/2021.08.04.455025

**Authors:** Katrin Geisler, Mark A. Scaife, Paweł M. Mordaka, Andre Holzer, Payam Mehrshahi, Gonzalo Mendoza Ochoa, Alison G. Smith

**Author notes:** Correspondence: Alison Smith.

## Abstract

*Chlamydomonas reinhardtii* has many attractive features for use as a model organism for both fundamental studies and as a biotechnological platform. Nonetheless, despite the many molecular tools and resources that have been developed, there are challenges for its successful engineering, in particular to obtain reproducible and high levels of transgene expression. Here we describe a synthetic biology approach to screen several hundred independent transformants using standardised parts to explore different parameters that might affect transgene expression. We focused on terminators and, using a standardised workflow and quantitative outputs, tested 9 different elements representing three different size classes of native terminators to determine their ability to support high level expression of a GFP reporter gene. We found that the optimal size reflected the median size of element found in the *C. reinhardtii* genome. The behaviour of the terminator parts was similar with different promoters, in different host strains and with different transgenes. This approach is applicable to the systematic testing of other genetic elements, facilitating comparison to determine optimal transgene design.

## 1. Introduction

Of the plethora of microalgae identified to date, only a small number have been studied extensively and even fewer have been developed sufficiently for their laboratory culture, biochemical analysis, and genetic manipulation to be considered routine [1]. Of these, the green alga *Chlamydomonas reinhardtii* (Chlorophyta) is the best studied, and over >30 years of basic and applied research, its study has contributed to our understanding of many fundamental biological processes, in particular photosynthesis [2] and motility [3]. It is often referred to as a model alga, and this designation has helped to galvanise efforts to develop several essential resources, which are now enabling the use of *C. reinhardtii* for biotechnology [4–6]. These include well-established basic laboratory protocols for culturing and molecular analysis, excellent genetics [7], fully sequenced genomes for the nucleus, mitochondrion and chloroplast [8], and efficient transformation protocols for all three genomes [9]. Routinely this is via electroporation [10], but simple vortexing with glass beads is also possible [11]. Such resources have led directly to the generation of an impressive array of genetic elements to enable transgene expression, including selectable markers, reporter genes [12–14] and promoter elements (reviewed in [4,5]), as well as gene silencing techniques that utilise RNA interference (RNAi) [15] or artificial microRNAs (amiRNAs) [16] and gene editing via CRISPR/Cpf1 [17]. In parallel, the implementation of synthetic biology approaches, most notably the adoption of standard parts and modular cloning methods, has enabled much more rapid generation of constructs for transformation and effective comparison between different designs, culminating in the generation of a kit of over 100 standardized elements [18], available at the Chlamydomonas Resource Center (https://www.chlamycollection.org/).

As a result, there have been many reports of the use of *C. reinhardtii* as a host for the production of high-value compounds and of individual proteins, particularly those with therapeutic potential (reviewed in [4,19,20]). The chloroplast has proved a versatile expression platform for recombinant proteins since it naturally accumulates high levels of soluble protein. *C. reinhardtii* is one of only two organisms, the other being *Nicotiana* spp., where chloroplast transformation is robust and routine. Transgenes can be introduced into the chloroplast genome by homologous recombination at precise locations using flanking homology arms bordering the gene(s) to be inserted. For several proteins, yields of up to 5% total soluble protein have been reported (reviewed in [21]). Disulphide bond formation occurs readily, but there is no glycosylation system in the organelle, so it is not suitable for expression of all proteins. In fact, more common is the introduction of transgenes into the nucleus, allowing a greater range of genes to be expressed, and the protein products can then be targeted to the appropriate subcellular location, including the chloroplast, by inclusion of appropriate targeting sequences in the genetic constructs [22]. Many of these are now available via the Chlamydomonas Spatial Interactome (https://sites.google.com/site/chlamyspatialinteractome/), which provides the means to target proteins not just to the organelles themselves, but also to specific sub-compartments within them [23]. As well as single proteins [24], there is increasing success in expressing metabolic enzymes to allow production of compounds such as the diterpenoids casbene, taxadiene, and 13*R*(+) manoyl oxide [25].

However, whilst it is straightforward to obtain large numbers of nuclear transformants of *C. reinhardtii*, transgenes are inserted essentially randomly. Ligation of trans-DNA is known to occur at sites of double-stranded breaks by the non-homologous end joining repair pathway. Insertion of the transgene is often preceded by endonucleolytic cleavage of DNA (with preference at CA/TG consensus motif), which results in the unexpected fragmentation and/or rearrangement of the transgene [26]. As a consequence, large numbers of transformants must be screened to ensure they contain the appropriate construct. More significantly, the level of transgene expression is often very low, with many transformants not expressing above background levels. Worse still, it is subject to gene silencing over time, particularly if selection is not maintained, most likely at the transcriptional level [27]. Although the reasons for this extensive silencing are not clear, several empirical approaches have been followed to try to overcome low or unreliable transgene expression (reviewed in [28]).

A particular milestone in the field of *C. reinhardtii* engineering was achieved as the result of UV mutagenesis of a strain that was only weakly expressing the CRY1 gene that confers resistance to emetine [29]. Two mutants, UVM4 and UVM11, were found that were resistant to much higher concentrations of emetine, and they were subsequently shown to express another, newly transformed, heterologous transgene at high frequency and to high levels. The common mutation between these two lines was subsequently shown to be a defect in the *SIR2* gene encoding a histone deacetylase [30].

As well as the genetic background of the host strain, design of constructs is also crucial. *C. reinhardtii* has very high GC content (with an average GC content of 64% throughout the nuclear genome and 68% in coding sequences [8]), and a strict biased codon usage [31]. Efficient transgene expression is achieved only by using appropriate codon usage, which in practice means that heterologous genes have to be synthesised. Inclusion of introns, which facilitate interaction of the RNA polymerase with the spliceosome, is similarly necessary, with an increase in expression of up to 5-fold observed over transgenes without introns in several studies (e.g. [32,33]). Finally, there have been extensive studies on optimal regulatory elements, particularly promoters. For example, there are several strong promoters that have been widely used including for the small subunit of Rubisco (RBCS) and a Photosystem I subunit (PSAD), but these are both preferentially expressed in the light [34] and *C. reinhardtii* does not appear to have strong constitutive promoters expressed under all conditions. Instead, researchers have developed novel promoters, such as the chimeric *HSP70A/RBCS2i* promoter [35] and several completely synthetic promoters based on combining sequence and nucleotypic characteristics of known promoters [36]. The choice of terminator has also been found to play a role. Kumar et al. [37] tested three different terminators in combination with seven different promoters. They found that maximal expression of the luciferase reporter gene was obtained using the *PSAD* promoter together with the *PSAD* terminator, although a *β2-tubulin* promoter / *PSAD* terminator combination was also effective, indicating that it was not simply cognate pairing that provided optimal expression levels. A further study identified terminators of highly expressed genes for a ribosomal subunit (*RPL23*) and ferredoxin (*FDX1*) as supporting high levels of luciferase reporter activity [38], and that the former was more effective than the *PSAD* terminator in expressing the *sh-ble* gene conferring zeocin-resistance. Here we extend this investigation of terminators further by analysing their size ranges in the *C. reinhardtii* genome and then test nine different terminators that represent the different size classes for their efficacy in regulation of transgene expression in the *C. reinhardtii* nucleus. We also present a standard workflow that has been developed in our laboratory for the effective nuclear transformation of *C. reinhardtii* and the quantitative measurement of several parameters to facilitate optimisation of construct design.

## 2. Materials and Methods

### 2.1. C. reinhardtii strain information

Three different *C. reinhardtii* strains were used in this work: the cell wall deficient strains CC-1615 cw15 mt- (cw15) (Chlamydomonas Resource Centre) and UVM4 [29], and the wild-type strain 12 (WT12), which is a walled strain derived from strain CC-124 137c (mt-nit1 nit2). *C. reinhardtii* cultures were grown in liquid or solid Tris-acetate phosphate (TAP) medium with Kropat’s trace elements [39], but excluding selenium at 25°C under continuous light conditions (light intensity of 80-100 µE·m^-2^·s^-1^).

### 2.2. Bioinformatic analysis

Version 5.6 of the *Chlamydomonas reinhardtii* (CC-503 v5.6) reference genome assembly [8] was downloaded from JGI’s plant genomics research database Phytozome 12 (https://phytozome.jgi.doe.gov/pz/portal.html) in March 2019. Filtered gene models ‘Creinhardtii_281_v5.6.gene.gff3’ were employed, together with custom R scripts, to determine genomic distance between subsequent genes and other genomic features, respectively. Annotations included coding regions of genes as well as position of individual 5’ and 3’ UTRs. Histograms were plotted applying a binsize of 100 bp.

Gene expression ranking was performed based on FPKM values of 17,824 nuclear, chloroplast or mitochondrial encoded genes obtained by RNA-seq of *C. reinhardtii* over a 24h period [34]. FPKM values from Strenkert et al. [34] were used to calculate mean expression levels, standard deviation as well as expression ranking. For expression over the diurnal cycle FPKM values from all timepoints were used.

### 2.3. Plasmid design and assembly

Constructs were made using parts encoding promoters, 5’ UTRs and terminators from *C. reinhardtii* genomic DNA as well as in-house plasmid templates for reporter genes and selectable markers generated by the Smith laboratory (University of Cambridge) or colleagues in other groups, as described in Tables S2 and S3. Genetic parts were amplified using primers listed in Table S1 and PCR products were purified with the Monarch PCR & DNA Clean up kit (NEB). Parts were assembled into plasmid constructs mainly by Gibson assembly [41] using the isothermal method, at 50 °C for 1 h. Alternatively, constructs were generated following standard Golden Gate (GG) cloning according to the modular cloning (MoClo) system [42,43], using parts from the Chlamydomonas MoClo toolkit [18] and parts that were created for this work. If necessary, parts were domesticated removing BpiI and BsaI sites by PCR-based mutagenesis and cloned into corresponding vectors by GG cloning using either BsaI (NEB) or BpiI (Thermo Fisher Scientific) depending on the level of the destination MoClo vectors, together with T4 Ligase (Thermo Fisher Scientific). Plasmids were propagated in *E. coli* (NEB 5-alpha competent cells), isolated using the Monarch Plasmid preparation kit (NEB) and verified by sequencing (Source Bioscience, Genewiz) or restriction digestion.

### 2.4. Chlamydomonas transformation and culturing

Plasmids were linearised and transformed into *C. reinhardtii* via electroporation as previously described [44]. Primary selection was performed on TAP agar plates supplemented with zeocin (10 µg ml^-1^) or paromomycin (10 mg l^−1^). Single colonies (primary transformants) were transferred to 96 well microtitre plates for culture in TAP liquid media, supplemented with antibiotics (5 mg l^−1^zeocin or 10 mg l^−1^ paromomycin). Following three sequential subcultures, each lasting seven days, a subset of viable cell lines were dubbed stable transformants and assessed for the expression of GFP via confocal microscopy. For analysis of the different terminators, promoters and host strains, each construct was tested in at least 3 independent transformations and the results presented are the average of >288 transformed lines.

### 2.5 Confocal Laser Scanning Microscopy

*C. reinhardtii* transformants carrying the Ble-GFP expression cassettes were grown for seven days in TAP media before visualization in a confocal laser scanning microscope (TCS SP5, Leica Microsystems, Germany). Images were acquired with excitation at 476 nm and emission detection between 485 and 518 nm for the GFP channel, and between 650 and 720 nm for the chlorophyll channel. Pictures were taken with a line average of 4 by the Leica LAS software.

### 2.6 Statistical tests

Transformation efficiencies were recorded as colonies appearing on selection plates after 7-10 days of growth for strains cw15 and UVM4, and 15-20 days for WT12 transformants. Algal growth in microtitre plates and GFP expression was assessed and scored after 7 days of growth. Values are given as means of at least three independent experiments ± standard error of the mean. Algal lines transformed with the *RBCS2* terminator construct were used for comparison unless otherwise stated and two-tailed Mann–Whitney U-tests were performed to determine statistical significance. A *p-*value of <0.05 was considered statistically significant.

## 3. Results

### 3.1 *Transgene instability: a recurrent problem in* C. reinhardtii

Transformation of a cell wall-deficient strain of *C. reinhardtii* such as cw15 by electroporation using an optimised protocol [10] will routinely yield >1000 transformants per µg of plasmid DNA within 7-10 days after plating on selective medium. However, only a proportion of these colonies will express the transgene effectively. We demonstrate this with results from a typical experiment as illustrated in Figure 1. *C. reinhardtii* cw15 cells were transformed with a construct encoding a Ble-GFP fusion protein under the control of the *HSP70<u>A</u>/<u>R</u>BCS2* chimeric promoter (*AR* promoter; [35]) and *RBCS2* 3’UTR/terminator, a combination that has been widely used in the literature [18]. The sh-*ble* gene from S*treptoalloteichus hindustanus* encodes a zeocin-binding protein, which is targeted to the nucleus, so that the green fluorescent protein (GFP) is also localises there [45]. Transformants were selected on TAP agar plates containing 10 µg/ml zeocin and then 96 colonies were picked into liquid medium (TAP+ 5 µg/ml zeocin) in a microtitre plate and subcultured every seven days. Cell growth was scored and recorded before every subculture. After three subcultures many of the initially picked lines were unable to grow under zeocin selection pressure (Figure 1, step 3). Those that were still viable were screened via confocal microscopy to identify lines that were expressing GFP (step 4). Some lines (e.g., line A3) showed high levels of fluorescence, whilst others (such as C12) had low level expression. Approximately 80% (e.g., line D4) did not express GFP above the detection limit. Thus, even when using a fusion protein, there was still variable expression of the GFP reporter in those that expressed sufficient Ble to be zeocin-resistant.

**Figure 1.**
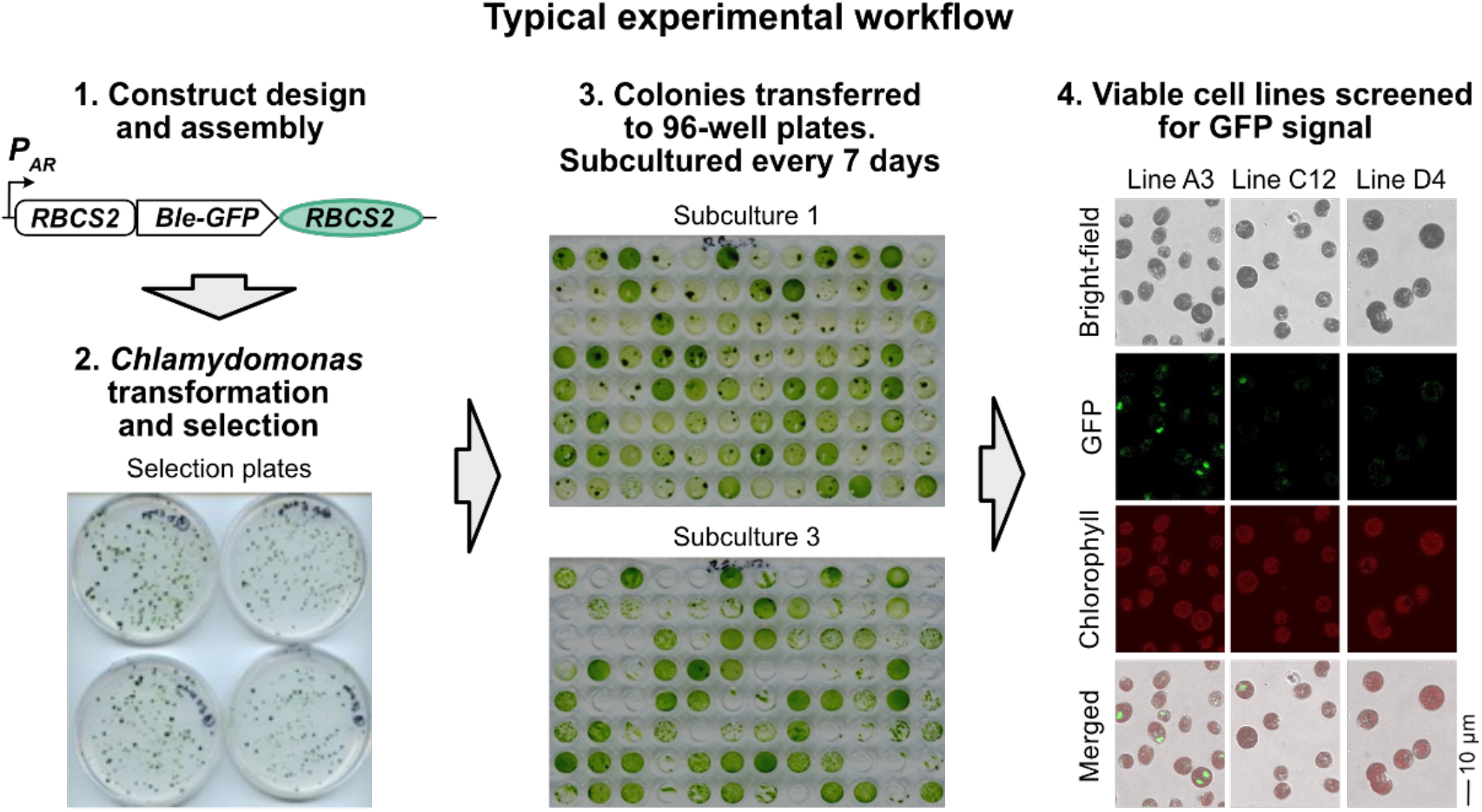
Typical experiment workflow of *C. reinhardtii* transformation and selection of stable transgenic lines. Step 1: Schematic of a DNA construct expressing a target transgene. A Ble-GFP reporter CDS is under the control of the *AR* promoter, and the *RBCS2* terminator. Step 2: After transformation of *C. reinhardtii* strain cw15 with a DNA construct, transformants are selected on agar plates containing zeocin. Step 3: Initially obtained colonies are transferred to TAP liquid media in microtitre plates, supplemented with zeocin, in a 96-well microtiter plate and grown in constant light, subculturing every seven days. The growth of the different cell lines is recorded. Step 4: After 3 subcultures, viable cell lines (ie zeocin-resistant) are screened for GFP expression via confocal microscopy. Representative images for three different cell lines (named according to the position in the 96-well plate) are shown using the brightfield, GFP and chlorophyll channels, as well as a merged figure of all three channels.

### 3.2 *Influence of terminators on transgene expression in* C. reinhardtii

Nonetheless, our workflow offered the means to screen large numbers of transformants to allow quantitative comparison between different constructs. We decided to explore in detail the impact of terminator choice on stability and level of gene expression since this has received less attention than other aspects of construct design [28]. Previous observations that using native features of the *C. reinhardtii* genome such as codon usage [31] and inclusion of introns [33,46] had a major impact on transgene expression, led us to reason that the same would be true for terminators. Accordingly, data from the *C. reinhardtii* genome (CC-503 v5.6) [8] were employed to assess the range and abundance of annotated features of all protein-coding genes, including the coding sequence and 5’ UTRs (Figure S1) and 3’ UTRs (Figure 2a). The annotated 3’ UTRs were delineated using transcriptome data sets [8] and include the canonical *C. reinhardtii* polyadenylation sequence of UGUAA [47,48] and so are a proxy for the terminator sequence. The size distribution graph shows that a sizeable number of genes (∼7,407; green bars in Figure 2a) have 3’ UTRs of between 100-600 bp. Around 11,000 genes have a predicted 3’ UTR of >600 bp (blue bars), and a smaller proportion (∼686 genes; orange bars) have short 3’ UTRs of <100 bp. Across the whole genome the median 3’ UTR length is 681 bp and the median distance between the 3’ UTR of one gene and 5’ UTR of the subsequent gene is 3,059 bp (Figure S1).

**Figure 2.**
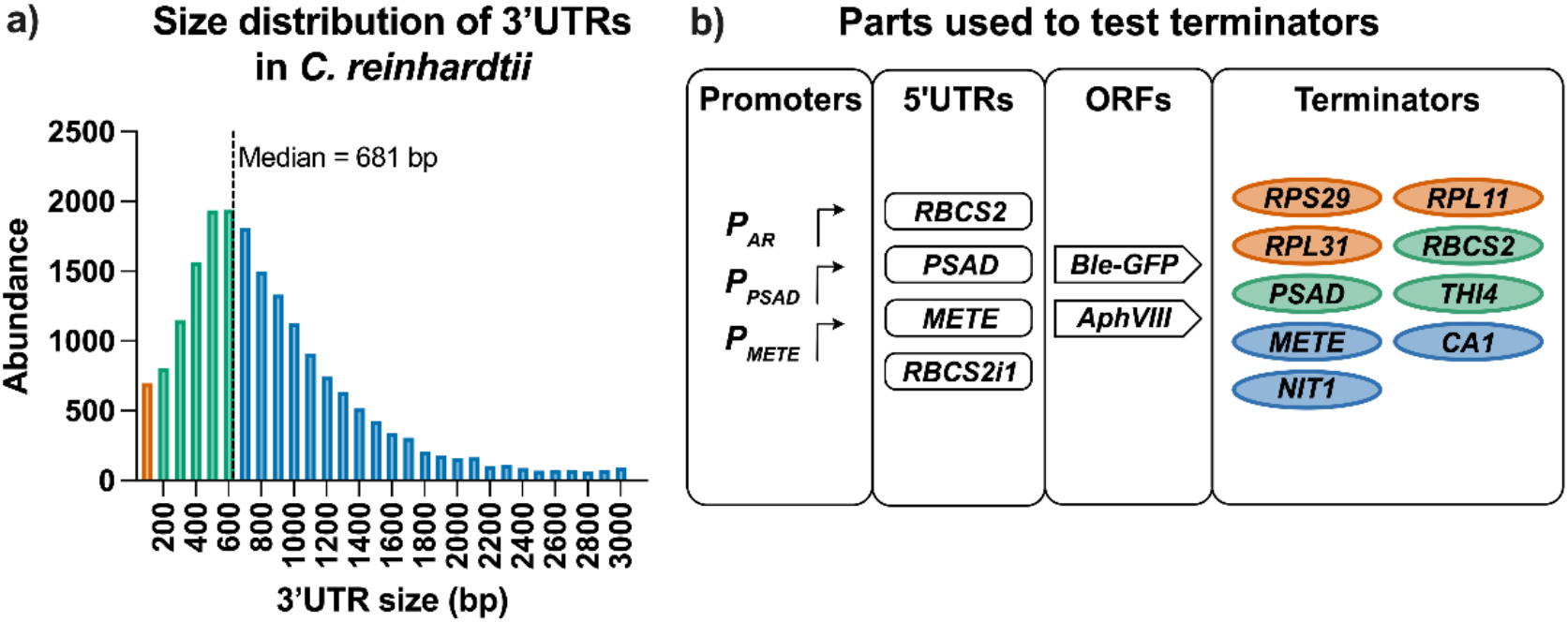
Analysis of 3’ UTR size and construct design to study terminators in *C. reinhardtii*. (a) Distribution of annotated 3’ UTR sizes from the *C. reinhardtii* genome v5.6 (b) Schematic of different promoter, 5’ UTR, ORF and terminator parts used throughout this work. Orange represents 3’ UTR/terminators of <100 basepairs (bp), green represents medium length 3’ UTRs (100–600 bp) and blue indicates long 3’ UTRs with a length > than the median. For details see Table 1. Promoter *P*_*AR*_ is a *HSP70/RBCS2* promoter fusion.

We took examples of these three different size classes of terminators to use as an independent variable in transgene expression using the workflow described above, testing their influence on transgene expression and stability. Three short elements chosen were ribosomal proteins, *RPS29, RPL11*, and *RPL31*, that were all under 100 bp. For those of medium length (150-500 bp), two (*RBCS2* and *PSAD)* have been widely used in *C. reinhardtii* constructs, and a third that is novel, *THI4*, encoding an enzyme of thiamine biosynthesis. The three representatives of the longer terminators with a length above 700 bp, were *METE* (encoding cobalamin-independent methionine synthase; [49]), *NIT1* (encoding nitrate reductase) and a carbonic anhydrase gene, *CA1* (Figure 2b, Table 1). We also include a no-3’UTR control that had no element beyond the stop codon. It should be noted that in previous studies [37] a 548 bp version of the *PSAD* terminator was used. As the annotated gene model predicted a shorter terminator, we opted to test a shorter version of 336 bp, which includes the UGUAA poly(A) signal. All other terminators used in this study also carry this UGUAA poly(A) motif.

**Table 1.**
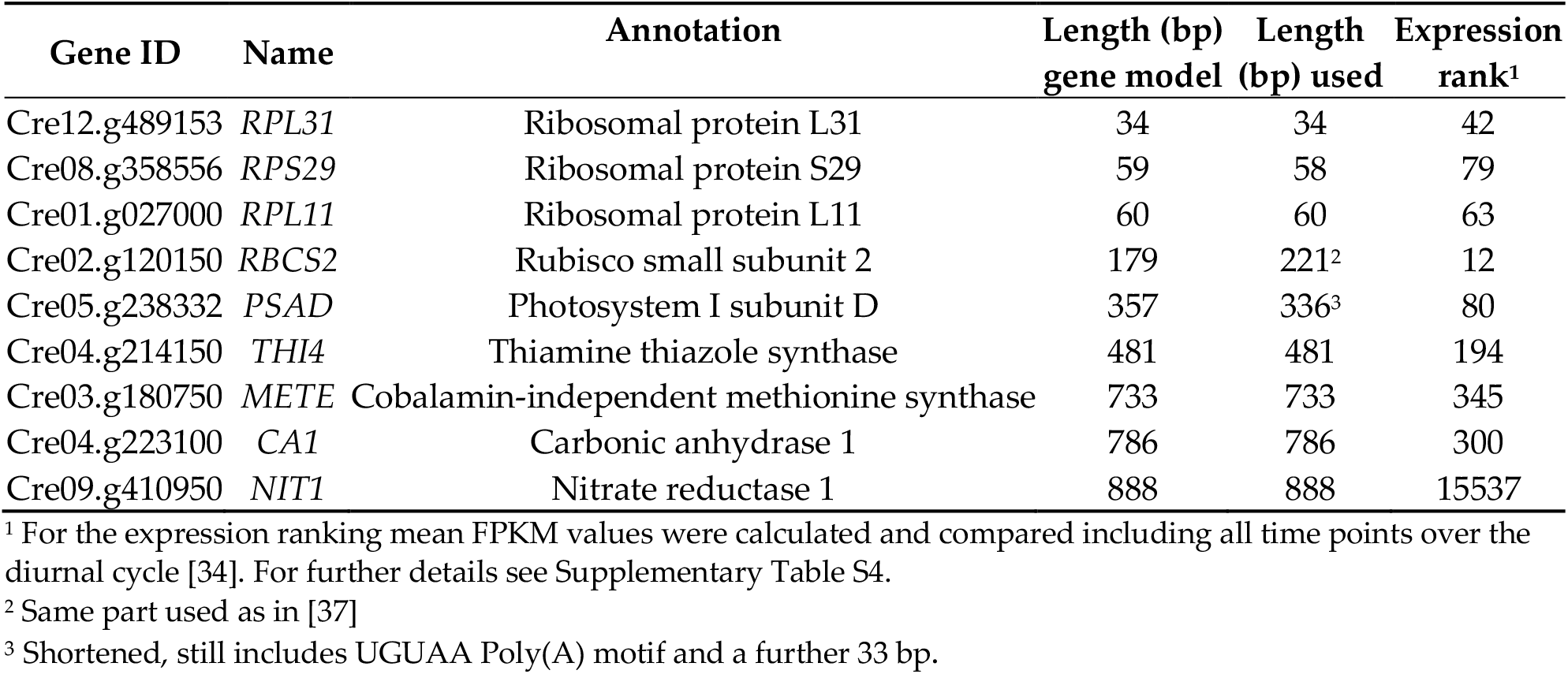
Candidate terminators tested in *C. reinhardtii*. The first three (*RPL31, RPS29, RPL11*) are short terminators (<100 bp), *RBCS2, PSAD and THI4* are classified as medium sized (100–600 bp), and *METE, CA1, NIT1* are long terminators (>600 bp). The predicted length based on the use of gene model analysis as well as the length of DNA parts used in this study is shown in base pairs (bp). The expression rank over the diurnal cycle was determined by comparing the mean FPKM values over all time points (from [34]).

| Gene ID | Name | Annotation | Length (bp)<br>gene model | Length (bp) used | Expression<br>rank<sup>1</sup> |
| --- | --- | --- | --- | --- | --- |
| Cre12.g489153 | *RPL31* | Ribosomal protein L31 | 34 | 34 | 42 |
| Cre08.g358556 | *RPS29* | Ribosomal protein S29 | 59 | 58 | 79 |
| Cre01.g027000 | *RPL11* | Ribosomal protein L11 | 60 | 60 | 63 |
| Cre02.g120150 | *RBCS2* | Rubisco small subunit 2 | 179 | 221<sup>2</sup> | 12 |
| Cre05.g238332 | *PSAD* | Photosystem I subunit D | 357 | 336<sup>3</sup> | 80 |
| Cre04.g214150 | *THI4* | Thiamine thiazole synthase | 481 | 481 | 194 |
| Cre03.g180750 | *METE* | Cobalamin-independent methionine synthase | 733 | 733 | 345 |
| Cre04.g223100 | *CA1* | Carbonic anhydrase 1 | 786 | 786 | 300 |
| Cre09.g410950 | *NIT1* | Nitrate reductase 1 | 888 | 888 | 15537 |

<sup>1</sup> For the expression ranking mean FPKM values were calculated and compared including all time points over the diurnal cycle [34]. For further details see Supplementary Table S4.
<sup>2</sup> Same part used as in [37]
<sup>3</sup> Shortened, still includes UGUAA Poly(A) motif and a further 33 bp.

In choosing these terminator elements, we considered their expression ranking by analysing previously published RNA-seq data of *C. reinhardtii* over a 24-hour period [34]. While there are diurnal expression differences in nearly every gene, the terminators chosen were from genes ranked highly (above rank 350) averaged over the entire diurnal cycle (Table 1, Table S1); terminators from highly expressed genes had previously shown to be effective in transgene expression [38]. The only exception is the *NIT1* terminator, whose average expression was ranked 15,537 in the analysed data set, but which is highly expressed in nitrogen deplete conditions [50].

In an initial experiment we assembled constructs in which Ble-GFP, containing the intron *RBCS2i1*, was under the control of the *AR* promoter and the *RBCS2* 5’ UTR, with each of the selected terminator candidates (Figure 2b). After plasmid linearisation, the different constructs were transformed into *C. reinhardtii* cw15 and transformants selected on TAP plates containing zeocin. The transformation efficiency differed considerably between them. With the *RBCS2* terminator (Figure 1) as a base line, we obtained ∼ 2.47 × 10^3^ colonies per µg DNA (Figure 3a). This is not statistically different from the number of colonies obtained when a no-3’UTR construct was used (∼ 1.63 × 10^3^ colonies per µg DNA). In contrast, the construct with the *CA1* terminator produced 5.50 × 10^3^ colonies per µg DNA, a 2.2-fold increase over the base case that was statistically significant (*p* <0.01). The *PSAD, THI4, METE* and *NIT1* terminators also increased the number of zeocin resistant transformants, but these were not statistically significant. None of the shorter terminators improved the transformation efficiency, despite being ranked in the top 100 most highly expressed genes in the diurnal RNA-seq dataset (Table 1; [34]). The trend suggests that the longer the terminator, the higher the number of initially selected transformants. We next investigated the stability of primary transformants through multiple rounds of subculturing under antibiotic selection pressure as described above (Figure 1). Figure 3b shows that the number of zeocin resistant cell lines decreased for all libraries over the course of this analysis. For our baseline construct carrying the *RBCS2* terminator, in three independent experiments an average of 74% of the initial lines were zeocin resistant after three subcultures (Figure 3c). The rate of decline in viability can be seen in the time course of Figure 3b, with the most rapid being in the no-3’UTR cell lines, with fewer than 50% surviving under constant selection pressure at subculture 3 (Figure 3c). All other tested terminators showed improved transgene stability, measured as zeocin-resistance, compared to the no-3’UTR library, retaining a level of viability above 65%. *CA1* demonstrated the greatest level of stability with a reduction in viability over the experimental period of just 13%. For the remaining seven terminators, a reduction in viability of between 20-32% was observed.

**Figure 3.**
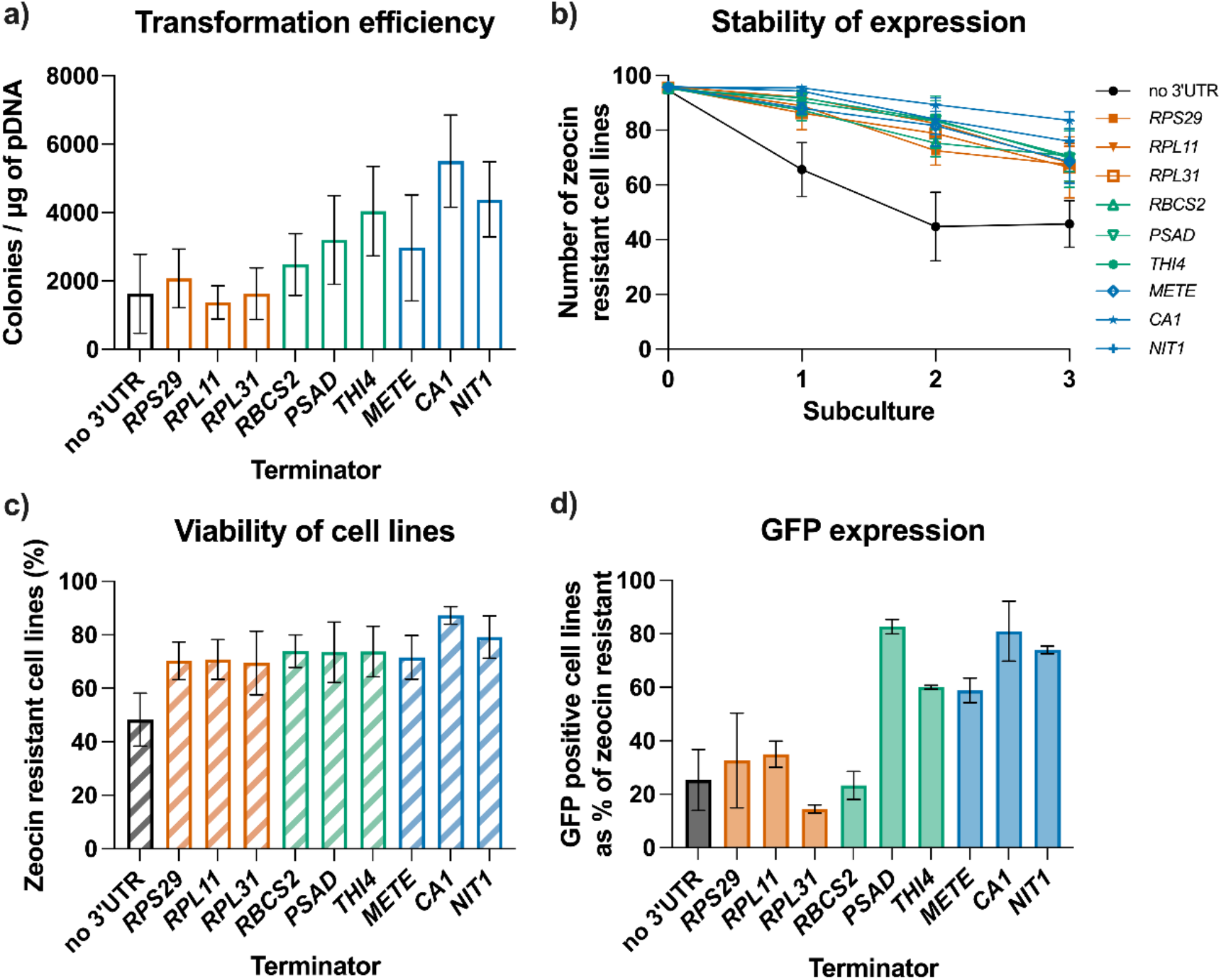
Analysis of the impact of terminators on transgene expression in *C. reinhardtii cw15*. *C. reinhardtii* was transformed with DNA constructs to test nine different terminators and a no-3’UTR control. (a) Transformation efficiency presented is the average of 3 replicate experiments, each containing a technical replicate (n=6). (b) Stability of transformants over 3 subcultures grown for seven days under constant selective pressure derived from three independent experiments, completed in duplicates, from which 96 independent transformants were analysed per experiment and construct. (c, d) Barplot of the percentage of (c) zeocin resistant cell lines at day seven of the third subculture and (d) barplot of the percentage of zeocin-resistant (ie viable) cell lines at day that were expressing GFP above the threshold measured by confocal microscopy. A subset of >30 individual cell lines per 3’ UTR were analysed. Error bars for all charts represent standard error of mean.

The stable cell lines were then examined via confocal microscopy to determine which were expressing detectable levels of GFP. As shown in Figure 3d, just 23% of the zeocin resistant cell lines carrying the RBCS2 terminator accumulated GFP (Figure 3d), comparable to that reported in other studies [31]. Similarly low percentages of GFP positive cell lines were observed for strains transformed with the short terminators of *RPS29, RPL11* and *RPL31* (15-35% GFP positive) or the no-3’UTR construct (25% GFP positive). The best performing constructs contained terminators from *PSAD*, with 83% GFP positives, and *CA1*, with 81% GFP positives, performing better than the *RBCS2* terminator (p <0.01). The *THI4, METE* and *NIT1* terminators also performed better than the shorter elements (*p <*0.05), with 60-70% GFP positives (Figure 3d).

### 3.3 *Activity of terminators in different* C. reinhardtii *host strains*

Host selection can have an influence on transgene expression. Cell wall deficient strains like the cw15 strain have been widely used to express transgenes, as they are more amenable to transformation compared to strains with an intact cell wall [51,52]. Strains UVM4 and its allelic variant UVM11, derived from a UV mutagenesis screen of a cw15 strain, exhibited enhanced expression of transgenes, accumulation of translated proteins and improved stable inheritance of the transgene over several generations [29]. However, cell wall-deficient strains have reduced motility and thus ability to mate compared to wildtype strains with an intact cell wall. In addition, upstream or downstream applications and limitations often make it necessary to use *C. reinhardtii* walled strains for the expression of transgenes since they are much less susceptible to shear or osmotic stress.

To determine whether the influence of terminators on transgene expression is dependent on strain selection, we tested a subset of the terminator constructs (no-3’UTR, *RPS29, PSAD* and *CA1*) in two other *C. reinhardtii* strains, wildtype 12 (WT12) and UVM4, as well as the previous used cw15. The same transformation protocol was used for all three strains. Colonies appeared on selection plates after 7 to 10 days for cw15 and UVM4 transformed cell and after 15 to 20 days for WT12 transformed cells. As expected, the overall transformation efficiency in the cell-walled strain WT12 was lower than the transformation efficiency for cw15 and UVM4. Nonetheless, the relative effects of the different terminators on transformation efficiencies were similar in the UVM4 and WT12 backgrounds as in cw15 (Figure 4a). Compared to the no-3’UTR control, the terminators of *PSAD* and *CA1* produced a statistically significant (p <0.01) higher number of zeocin selected transformants in each of the tested background strains. The *RPS29* terminator also gave more transformants compared to the no-3’UTR control, although the difference was not statistically significant (p >0.05).

**Figure 4.**
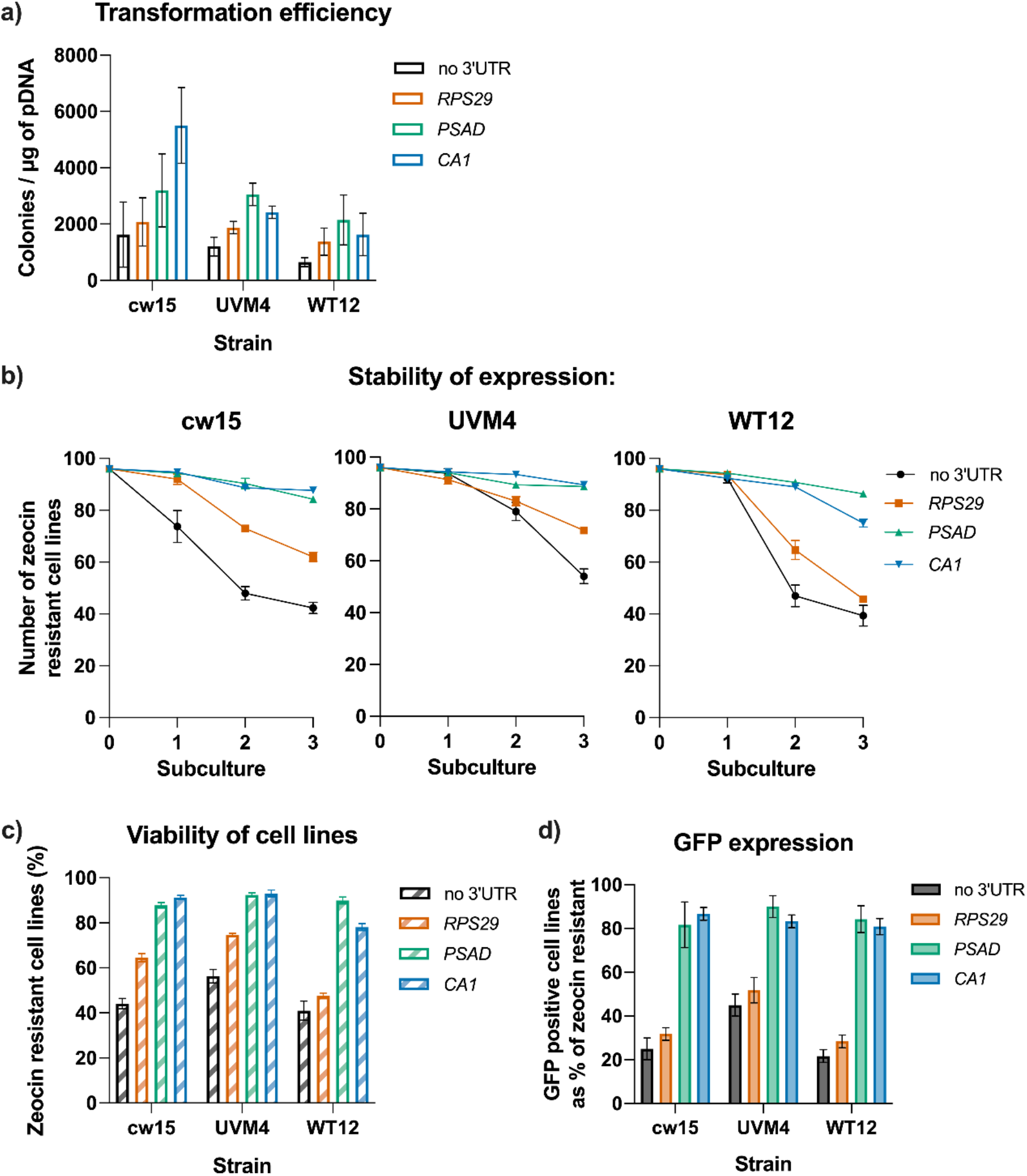
Influence of terminators on transgene expression in different *C. reinhardtii* strains. (a) Transformation efficiency for constructs carrying three different terminators (*RPS29, PSAD, CA1*) transformed into different *C. reinhardtii* strains: the cell wall deficient strains cw15 and UVM4, and the walled strain WT12. Presented are the averages of three replicate experiments. (b) Stability of *ble*-gene expression over three subcultures under constant antibiotic selection is shown for cw15 (left), UVM4 (middle) and WT12 (right panel). (c, d) Percentage of zeocin-resistant cell lines at day seven of the third subculture and (d) percentage of those zeocin resistant cell lines expressing GFP above the threshold detectable by confocal microscopy. Error bars for all charts represent standard error of mean. Data for strain cw15 is also shown in Figure 3.

A similar pattern in transgene stability between the different terminators was seen across the three host strains (Figure 4b and 4c). The decline in zeocin-resistant strains was most pronounced in the no-3’UTR and *RSP29* cell lines, whereas the *CA1* and *PSAD* lines retained a level of zeocin resistance of over 75% after 3 subcultures. Consistent with a previous report (Neupert et al., 2009), we observed that transgenes expressed in the UVM4 strain showed a higher stability, measured as resistance to zeocin, compared to the same constructs expressed in a WT12 or cw15 strain. For three of the four cell lines (*no-3’UTR, RPS29* and *CA1*), this higher transgene stability is statistically significant between strains (p <0.05) (Figure 4b and 4c).

Analysing the stable zeocin-resistant cell lines for GFP expression, again we observed the same trend across the three different strains. The percentage of cell lines expressing GFP is lower for strains transformed with constructs carrying no-3’UTR or the short *RPS29* element in all three background strains, than for those using the medium or long terminators, *PSAD* and *CA1* respectively (p <0.01) (Figure 4d). In all three background strains over 80% of the viable cells expressed GFP when either a *PSAD* or *CA1* terminator was used, with no statistical differences between them. In contrast, with the short *RPS29* terminator, the percentage of GFP positive cell lines dropped to 52% in the UVM4 background and to 32% and 28% in cw15 and WT12, respectively. Compared to the no-3’UTR control, the short RPS29 terminator has no significant influence on the percentage of GFP positive lines (p >0.05). This further reinforces the point that the use of medium and long terminators is preferred for stable and high transgene expression, independent of the *C. reinhardtii* host strain.

### 3.4 Analysis of the effect of promoter choice on transgene expression

To examine the impact of different promoter / terminator combinations on recombinant protein production, the promoter and 5’ UTR of *PSAD* and *METE* were employed in combination with the terminators of *CA1* and *PSAD* and compared with the original AR promoter (Figure 5). *PSAD* is considered a strong promoter, although it shows a bias for expression during the light period compared to the dark (Table S4; [34]), and *METE* is repressed by the addition of vitamin B12 and so has the potential to be a regulatory genetic element for biotechnology purposes (Helliwell et al., 2014). Plasmids were constructed, linearised and transformed into *C. reinhardtii* as described previously. Constructs using the *PSAD* promoter with either the *CA1* or *PSAD* terminator increased transformation efficiency 1.4 and 2.8-fold (*p* <0.05) respectively, when compared to the corresponding *AR* promoter / terminator line (Figure 5a). In contrast, those with the *METE* promoter showed reduced transformation efficiency compared to *AR*, to just 17 and 39% with *CA1* and *PSAD* terminators, respectively (*p* <0.05).

**Figure 5.**
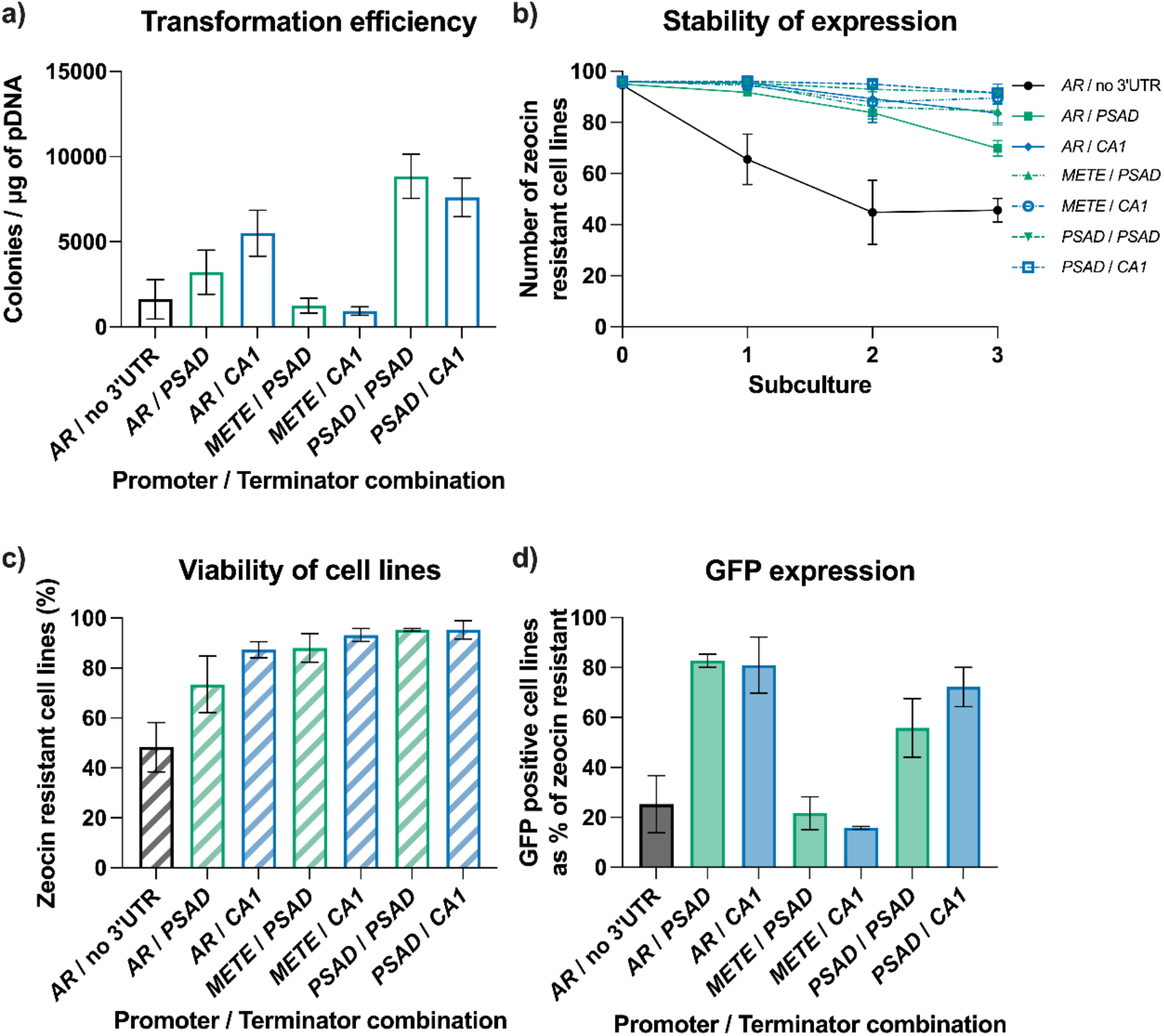
Analysis of the impact of different promoter and terminator combinations on transgene expression. The cell wall deficient strain cw15 was transformed with DNA construct expressing the Ble-GFP reporter under the control of different promoter / 5’ UTR and terminator combinations. The three promoters *AR, METE* and *PSAD* were combined with the *CA1* or *PSAD* terminator. Data from lines transformed with a construct carrying the AR promoter and no-3’UTR, shown in Figure 3, are also shown as reference. (a) Transformation efficiency for transformants carrying the different promoter / terminator combinations. Presented are the averages of three replicate experiments. (b) Expression stability of transformants over three subcultures under constant antibiotic selection. (c, d) Percentage of zeocin resistant cell lines at day seven of the third subculture that were also GFP-positive as determined by confocal microscopy. Error bars for all charts represent standard error of mean.

Nonetheless, both the *PSAD* and *METE* promoter cell lines, when employed with the terminator of *PSAD*, exhibited respectively 95% and 88% stability after three subcultures (Figure 5b and 5c). Comparable constructs that employed the *CA1* terminator showed a similar stability with both the *PSAD* and *METE* promoters. In the lines transformed with the *AR* promoter / no-3’UTR construct only ∼45% of initial picked cell lines remained zeocin resistant. Compared to this control, the percentage of zeocin viable cell lines is significant higher in all other tested promoter / terminator combinations (*p* <0.05).

To further investigate the influence of the promoter on average gene expression across a range of stable transformants confocal microscopy was used to test for GFP expression. When the promoter was changed from *AR* to *PSAD* no significant change in the frequency of GFP positives in the sample set was observed when the *CA1* terminator was used, although there was a significant reduction with the *PSAD* terminator, from 83% to 55% (p <0.05) (Figure 5d). Changing the *AR* promoter for the *METE* promoter produced an even greater reduction in the frequency of GFP positives in the sample, down to 16% with the *CA1* element (*p* <0.0005) and 22%, with the *PSAD* terminator (*p* <0.0005) (Figure 5d). These results might reflect the fact that the *METE* promoter element used here is noticeably weaker when controlling transgenes, possibly because other elements of the gene are necessary for high level expression [49,53].

### 3.5 Testing promoter and terminator combinations to drive expression of an independent transgene

So far, the analysis of the terminators has been with a single transgene expressing the fusion protein Ble-GFP, which was both the selection marker and reporter protein. We wanted to examine their influence on transgene expression *per se*, by including a different selection marker, in this case the paromomycin resistance gene *AphVIII*. The test expression cassette used the *Ble-GFP* transgene as before, under the control of the *PSAD* promoter and the three terminators used earlier, *RPS29, PSAD* and *CA1* (Figure 6a). This mimics metabolic engineering applications, where cassettes for introduced biosynthetic enzymes would be used together with a separate selection marker [44,54,55]. To allow the combinatorial assembly of different promoter / terminator combinations Golden Gate assembly following the MoClo syntax was applied [18]. We also included the intron *RBCS2i1* upstream of the gene coding regions to enhance transgene expression [13].

**Figure 6.**
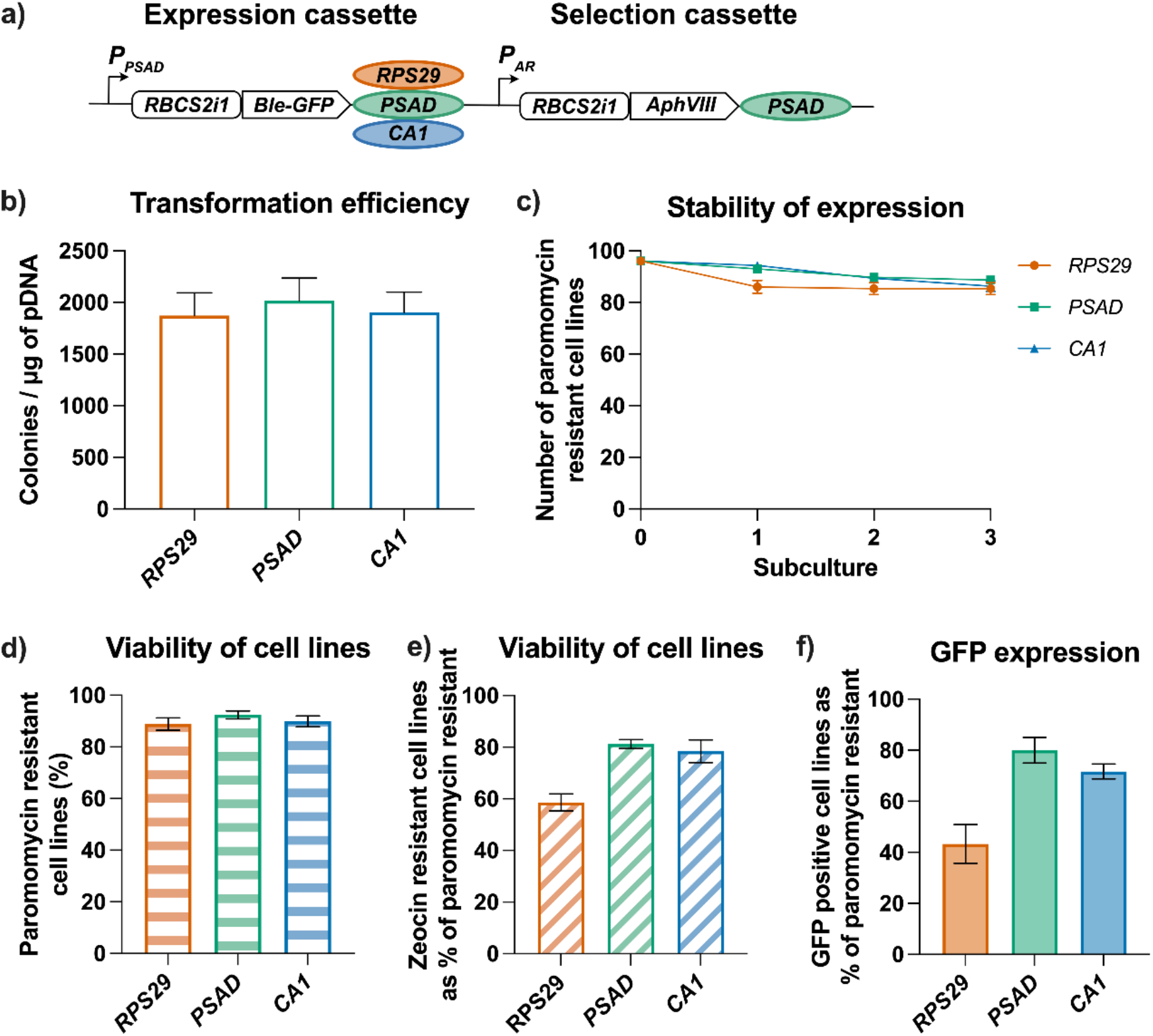
Testing terminators driving expression of transgenes in two different expression cassettes. (a) Schematic of construct design using two expression cassettes. The selection cassette is driving expression of the paromomycin resistance gene *AphVIII* using an *AR* promoter / *PSAD* terminator combination. In the Ble-GFP expression cassette the *PSAD* promoter is driving expression and three different terminators are tested: *RPS29, PSAD* and *CA1*. Constructs were introduced into strain UVM4. (b) transformation efficiency for the average of 3 replicate experiments, each containing a technical replicate (n=6). (c) Stability of transformants over 3 subcultures under constant antibiotic selection of paromomycin. (d) Barplot of the percentage of paromomycin resistant (left), zeocin resistant (middle) and GFP positive cell lines (right panel) derived from the experiment.

UVM4 was transformed with the different constructs and transformants were selected on plates containing paromomycin. As expected, no difference in transformation efficiencies was observed with any construct (Figure 6b) because the selection was based on the identical cassette. Similarly, subculturing the primary transformants in TAP media with paromomycin and scoring growth to determine levels of *AphVIII* transgene expression, showed excellent stability over time with the *PSAD* terminator, with over 85% of the initial cells still paromomycin resistant (Figure 6c and 6d). It should be noted that the *PSAD* terminator used in the initial experiments (Figure 3-6) is 366 bp long, while the *PSAD* terminator L0 part from the MoClo kit is 548 bp long similar to the one used in previous studies [37].

After the third subculture, the paromomycin resistant cell lines were analysed for expression of the *Ble-GFP* transgene from the independent expression cassette, scoring both zeocin resistance and GFP expression. When the short terminator *RPS29* was used 59% of the tested cell lines were zeocin resistant and 43% expressed GFP above detectable limits of the confocal microscope (Figure 6e and 6f). The percentage is significantly increased when using the *PSAD* 3’ UTR (81% and 80%) or the *CA1* 3’ UTR (78% and 72%) for zeocin resistance or GFP fluorescence respectively (p <0.005). Again, this demonstrates that the longer terminators *PSAD* and *CA1* are more effective than shorter ones.

## 4. Discussion

In the work presented here we used a standardised workflow to demonstrate quantitatively the influence of 9 different terminators on transgene expression in *C. reinhardtii*. We found that the length of the terminator was the main factor in efficacy – the short terminators (<100 bp) of the three ribosomal subunits we tested, RPL11, RPL31 and RPS29, performed much less well in terms of transformation efficiency, transgene stability and level of expression than did the longer terminators (Figure 3), and were actually no better in terms of high level expression of GFP than the negative control (no-3’UTR), which had no defined sequence after the stop codon. This is despite the fact that the ribosomal subunits are more highly expressed across the diurnal cycle (Table 1; [34]) than the other genes. Similarly, we found that the 221 bp terminator element from the *RBCS2* gene, whose expression levels are ranked even higher than the ribosomal genes, was less effective, particularly in terms of expression level of the GFP reporter, than the other 5 elements (with lengths from 336 to 888 bp), even though all transgenes were driven by the chimeric *HSP70<u>A</u>/<u>R</u>BCS2* promoter (*AR*). Conversely, the *NIT1* terminator was effective in driving reporter gene levels above the threshold in the majority of stable transformants, despite its low expression ranking under the conditions of cell growth (in media containing ammonium ions and no nitrate). This lack of correlation between expression level of the native gene and the ability of its regulatory elements to confer similar properties on a transgene is consistent with the earlier observation by López-Paz et al. [38] on two other ribosomal proteins: the *RPL23* terminator was effective in promoting high level expression, but that for *RPL35a* was not, whilst their expression levels were both in the top rank in the diurnal RNA-seq dataset [56].

The terminators from *PSAD* and *CA1* gave the best results in all experiments. The effectiveness of the *PSAD* terminator for transgene expression in *C. reinhardtii* has been noted previously [18,37,38] although the reason for this is unknown. Our best understanding of the role(s) of terminators comes from studies in mammals and yeast, but similar features are known to operate in plants [57]. Terminators are crucial for transcription, where interaction between the 5’ and 3’ ends of the mRNA facilitate recycling of RNA polymerase II from the end of the gene to the start, stimulating initiation of transcription [58]. In addition, their interaction with the polyadenylation complex is essential for appropriate 3’end formation including addition of the poly(A) sequence [59], essential for stabilisation of the mRNA, export to the cytoplasm and efficient translation. Detailed dissection of plant terminator sequences has revealed that in addition to a polyadenylation motif of AAUAAA or AAUAAA-like, which is 10-20 nt from the cleavage site, there are several conserved near- and far-upstream elements (NUEs and FUEs respectively). The importance of these elements was confirmed by a study in maize where several ubiquitin terminators were analysed for their ability to drive reporter gene expression [60]. In contrast, a genome-wide analysis of the poly(A) sites in *C. reinhardtii* indicated that the conserved UGUAA motif (which is itself distinct from that in plants and animals) is the major *cis*-element involved in polyadenylation, with no other conserved sequences identified [48]. Since all the terminators we tested contained this sequence, this cannot be the reason for the differences we observed. It should be mentioned that there are many terminators in the *C. reinhardtii* genome much longer than the ones we studied here (Figure 2a) and which may contain additional regulatory sequences.

Whatever the explanation, the behaviours observed in the first experiment (Figure 3) were recapitulated when a subset of the terminators were tested in different host strains (Figure 4). Transformants with the *PSAD* and *CA1* terminators showed stable and high-level expression irrespective of the host, whereas *RPS29* performed the same or only slightly better than the no-3’UTR negative control. Stability of the zeocin-resistance phenotype in transformants was better in strain UVM4 than cw15 or WT12 as expected [29], particularly for those carrying the *PSAD* and *CA1* terminators (Figure 4b), but there was no discernible difference in the proportion of transformants expressing GFP (Figure 4d). However, it should be noted that these data are not the absolute level of GFP expression, simply a yes/no indication of whether or not fluorescence was observed by confocal microscopy. Exchanging the *AR* promoter for that from *PSAD* again demonstrated that the *PSAD* and *CA1* terminators conferred much better stability and expression of the transgene than the no-3’UTR negative control (Figure 5). When the *METE* promoter was used only an increase in stability was observed, with little or no GFP expression above baseline. This is likely to reflect the fact that *METE* promoter element used here is much weaker than *PSAD* or *AR* [49], so that whilst there is sufficient Ble protein to confer zeocin-resistance, GFP fluorescence is at or below the minimum detection threshold. When the *RPS29, PSAD* and *CA1* terminators were used to regulate the transgene separately from the selectable marker they again behaved as predicted (Figure 6). Together, these results indicate that the terminator parts are acting in an orthogonal manner, meaning that they could be expected to behave similarly in other construct design. Indeed, we have successfully used the *CA1* terminator to express casbene synthase from the higher plant *Jatropha curcas* in *C. reinhardtii* [44]; the engineered strain is still producing casbene at the same level over 2 years later (data not shown).

Our aim in this work has been to enhance the already extensive synthetic biology toolkit for *C. reinhardtii* to increase the flexibility for transgene expression in this organism. The advantage of the approach we have taken here to screen for optimal genetic parts, in this case terminator elements, is that with a defined workflow, we were able to explore the parameter space across a range of different characteristics systematically and quantitatively. The method is relatively rapid (4-5 weeks from transformation to output on the activity of the parts). To facilitate visualisation of the results that we collected and allow an objective assessment of which would be the most suitable to use in subsequent construct designs, we combined them in Figure 7, plotting the stability of the zeocin-resistance against the proportion of the stable lines expressing GFP above the threshold. The size of the circles represents the transformation efficiency. In Figure 7a, all the data from cw15 are presented, whereas in Figure 7b, the data available from all 3 strains are shown. There is a clear trend of a positive correlation between the three parameters that are quantified, with the constructs containing the *PSAD* and *CA1* terminators clustering together in the top right of the graphs. Thus even though differences in the measured parameters are relatively small, combining the measurements provides more confidence in the choice of optimal parts for any particular application.

**Figure 7.**
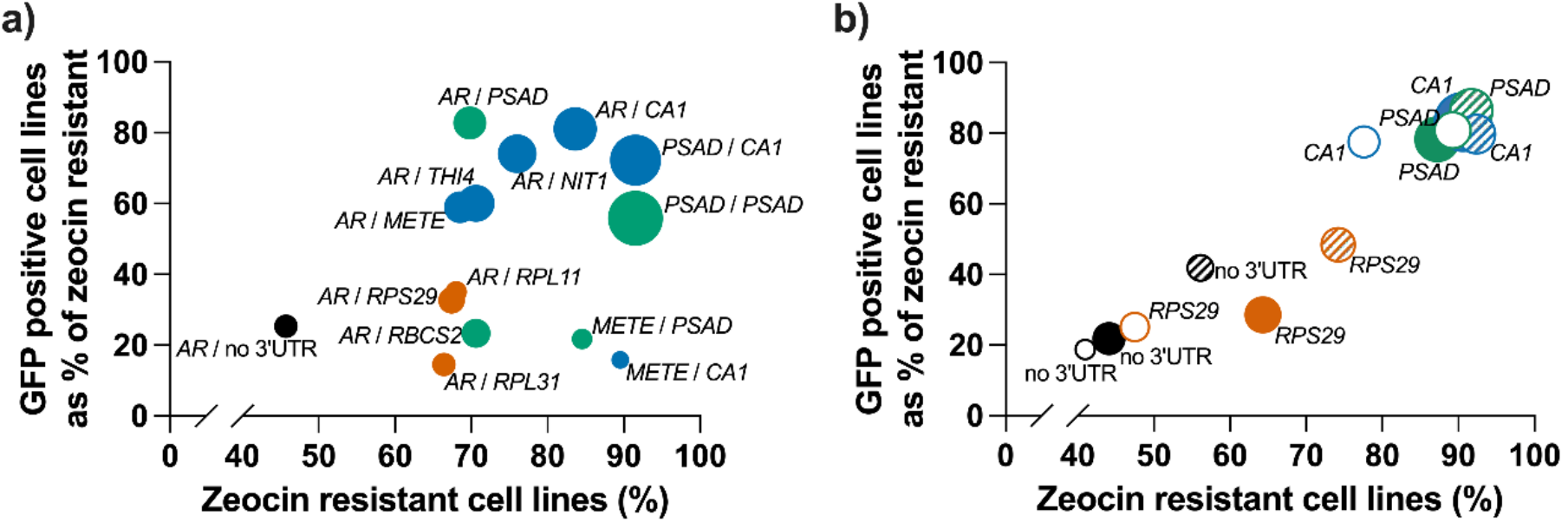
Impact of terminators on different parameters to measure transgene expression in *C. reinhardtii*. (a) The influence of different promoter / terminator combinations on transformation efficiency (circle size), resistance to zeocin (x-axis) and GFP expression (y-axis) tested in *C. reinhardtii* cw15. (b) The effect of a subset of terminators (*RPS29, PSAD, CA1*) on transformation efficiency (circle size), resistance to zeocin (x-axis) and GFP expression over the detection limit (y-axis) when tested in different *C. reinhardtii* strains, cw15 transformants (filled circles), UVM4 transformants (hatched circles) and WT12 transformants (open circles). Shown are average data from three independent transformation experiments.

In summary, we have expanded the set of terminators that can be used for stable and high transgene expression in *C. reinhardtii*. The *CA1* terminator behaves very similarly to the well-characterised *PSAD*, and so offers the means to use distinct but effective terminators for the selectable marker and the transgene in a typical construct design. Both of these are part of the original MoClo kit [18], which enables rapid assembly via modular cloning. In addition, three other terminators, from the *NIT1, METE* and *THI4* genes, have been shown to be effective, and they too have been domesticated for use in the same system and syntax. All are available from the Chlamydomonas Resource Centre.

## Supporting information

Supplementary meterial

## Supplementary Materials

The following are available online.

Figure S1: Analysis of genomic features in *C. reinhardtii*., Table S1: Primers used for Gibson assembly and DNA part generation., Table S2: List of plasmids generated by Gibson assembly, Table S3: MoClo constructs employed and generated, Table S4: Expression ranking of *C. reinhardtii* genes used in this study, Dataset S1: Distribution of genomic elements in *C. reinhardtii* genome, Dataset S2: Gene expression values from Strenkert et al.

## Author Contributions

Conceptualization, A.G.S.; methodology, K.G., M.A.S. and A.H.; formal analysis, K.G., M.A.S. and A.H.; investigation, K.G., M.A.S., A.H. and P.Me.; data curation, K.G., M.A.S. and A.H.; writing— original draft preparation, K.G.; writing—review and editing, K.G., P.Me., P.Mo., G.M.O., A.H. and A.G.S.; visualization, A.H. and P.Mo.; supervision, A.G.S.; project administration, A.G.S; funding acquisition, A.G.S. All authors have read and agreed to the published version of the manuscript.

## Data Availability Statement

The data is available in the data repository of the University of Cambridge, UK.

## Acknowledgments

We are grateful for laboratory support from Eleanor Tomsett, who helped with data collection, Faye Loo, who helped with cloning of DNA constructs, Fiona Taylor and Katie Sutherland for algal subculturing and Sue Aspinall and Lorraine Archer for laboratory management. *C. reinhardtii* strain UVM4 was obtained from Prof. Dr. Ralph Bock, MPI-MP, Golm, Germany.

