## Supplementary meterial for "Exploring the impact of terminators on transgene expression in *Chlamydomonas reinhardtii* with a synthetic biology approach"

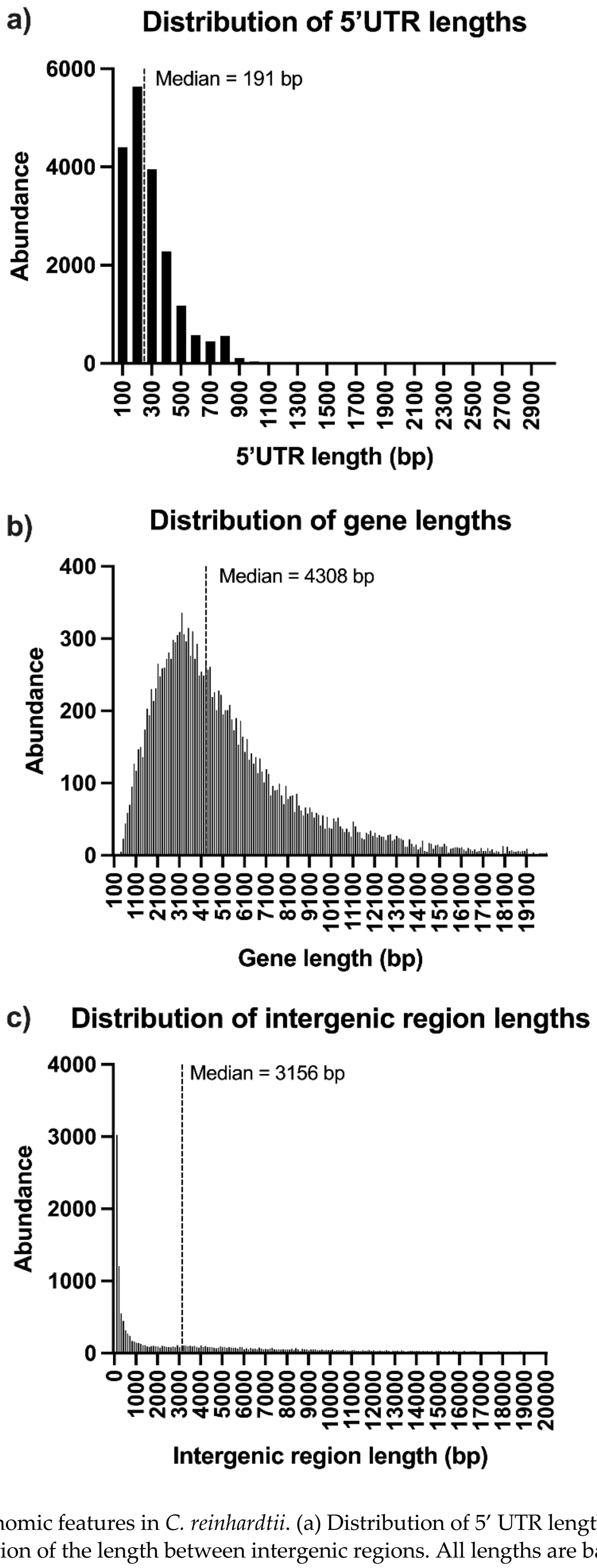


**Figure S1.** Analysis of genomic features in *C. reinhardtii*. (a) Distribution of 5′ UTR lengths. (b) Distribution of gene lengths. (c) Distribution of the length between intergenic regions. All lengths are based on annotations from the *C. reinhardtii* genome v5.6. Calculated median sizes of genomic features are shown.

**Table 1.** Primers used for Gibson assembly and DNA part generation.

| Part Name | Primer Name | Primer Sequence (5′-3′) |
|---|---|---|
| GFP-3′UTR | Rbcs2(B)-Ble F | GAGAAGTCACTCAACATCTTAAATGGCCAAGCTGACCAGCGCCGTTC |
| GFP-RPL31 | GFP-L31 R | CCACAGAAAGAGTTACACACTTGCGGCGTATGGCTTACTTGTACAGCTCGTCCATGCCGT |
| GFP-RPS29 | GFP-RPS29 R | ACGGCATGGACGAGCTGTACAAGTAAAATGCCCGAATGTTGGGTATCTAGCTCACACG CAGTTTGTAAGGTGCTGAGGCGTTG |
| GFP-RPL11 | GFP-RPL11 R | GGTCAGGACTTGTTACACAGATGGGGTGGCATCTGTCATCACCAGCCTCTTGTGGCCGCT TTACTTGTACAGCTCGTCCATGCCGT |
| GFP-RBCS2 | GFP-RBCS2 R | GCTCAGATCAACGAGCGCCTCCATTTACTTGTACAGCTCGTCCATGCCGT |
| GFP-PSAD | GFP-PSAD R | GGTACAGGCGGTCCAGCTGCTGCCTTACTTGTACAGCTCGTCCATGCCGT |
| GFP-THI4 | GFP-THI4 R | GCAGCCCAGTAGGTTCTGAGCGCGCTTACTTGTACAGCTCGTCCATGCCGT |
| GFP-METE | GFP-METE R | TCAGCATAAATCAAGGCAGGCAGCTTACTTGTACAGCTCGTCCATGCCGT |
| GFP-CA1 | GFP-CA1 R | TAGCGTGACTAACTACTGGGAAGTTTACTTGTACAGCTCGTCCATGCCGT |
| GFP-NIT1 | GFP-NIT1 R | ACCGTGGCCACAATCTCTAGCGATTTACTTGTACAGCTCGTCCATGCCGT |
| GFP-noUTR | GFP-noUTR R | GTGCGTCGGGTGATGCTGCCAACTTACTGATTTAGTTACTTGTACAGCTCGTCCATGCCGT |
| noUTR | GFP-noUTR F | ACGGCATGGACGAGCTGTACAAGTAACTAAATCAGTAAGTTGGCAG |
| RBCS2 | GFP-RBCS2 F | ACGGCATGGACGAGCTGTACAAGTAAATGGAGGCGCTCGTTGATCTGAGC |
| RBCS2 | RBCS2-BADCmR | GTGATGCTGCCAACTTACTGATTTAGCGCTTCAAATACGCCCAGCCCGCC |
| PSAD | GFP-PSAD F | ACGGCATGGACGAGCTGTACAAGTAAGGCAGCAGCTGGACCGCCTGTACC |
| PSAD | PSAD-BADCmR | GTGATGCTGCCAACTTACTGATTTAGCACGGCAAACTCTCACATGGCCTG |
| THI4 | GFP-THI4 F | ACGGCATGGACGAGCTGTACAAGTAAGCGCGCTCAGAACCTACTGGGCTGC |
| THI4 | THI4-BADCmR | GTGATGCTGCCAACTTACTGATTTAGTCTCAACTCCAAATTGTTACATGA |
| METE | GFP-METE F | ACGGCATGGACGAGCTGTACAAGTAAGCTGCCTGCCTTGATTTATGCTGA |
| METE | METE-BADCmR | GTGATGCTGCCAACTTACTGATTTAGACGTCACACCTTGCTACCCTACAG |
| CA1 | GFP-CA1 F | ACGGCATGGACGAGCTGTACAAGTAAACTTCCCAGTAGTTAGTCACGCTA |
| CA1 | CA1-BADCmR | GTGATGCTGCCAACTTACTGATTTAGTCCATGGGCATCTTACATGTCGTA |
| NIT1 | GFP-NIT1 F | ACGGCATGGACGAGCTGTACAAGTAAATCGCTAGAGATTGTGGCCACGGT |
| NIT1 | NIT1-BADCmR | GTGATGCTGCCAACTTACTGATTTAGCGCTTGCTTAATTTACAACTGTGC |
| RPL31-pUC | L31-BADCmF | GCAAGTGTGTAACTCTTTCTGTGGCTAAATCAGTAAGTTGGCAGCATCAC |
| RPS29-pUC | RPS29-BADCmF | AATGCCCGAATGTTGGGTATCTAGCTCACACGGCAGTTTGTAAGGTGCTGAGGCGTTGCT AAATCAGTAAGTTGGCAGCATCAC |
| RPL11-pUC | RPL11-BADCmF | AGCGGCCACAAGAGGCTGGTGATGACAGATGCCACCCCATCTGTGTAACAAGTCCTGAC CCTAAATCAGTAAGTTGGCAGCATCAC |
| RBCS2-pUC | | GGCGGGCTGGGCGTATTTGAAGCGCTAAATCAGTAAGTTGGCAGCATCAC |
| PSAD-pUC | PSAD-BADCmF | CAGGCCATGTGAGAGTTTGCCGTGCTAAATCAGTAAGTTGGCAGCATCAC |
| THI4-pUC | THI4-BADCmF | TCATGTAACAATTTGGAGTTGAGACTAAATCAGTAAGTTGGCAGCATCAC |
| METE-pUC | METE-BADCmF | CTGTAGGGTAGCAAGGTGTGACGTCTAAATCAGTAAGTTGGCAGCATCAC |
| CA1-pUC | CA1-BADCmF | TACGACATGTAAGATGCCCATGGACTAAATCAGTAAGTTGGCAGCATCAC |
| NIT1-pUC | NIT1-BADCmF | GCACAGTTGTAAATTAAGCAAGCGCTAAATCAGTAAGTTGGCAGCATCAC |
| pUC-AR | pUC-AR R | CACCAATCATGTCAAGCCTCAGCGAGCTCCCCGCCGTCGTACCGAGCTCGAATTCGTAAT CATGGTCA |
| pUC-PSAD | pUC-PSAD R | TGACCATGATTACGAATTCGAGCTCGGTACATCCCACACACCTGCCCGTCTGCCTGACA |
| pUC-METE | pUC-METE R | GATTACGAATTCGAGCTCGGTACGTAGGTCAGGACCAGAGCCTACAAC |
| AR prom | pUC-AR F | CGAATTCGAGCTCGGTACGACGGCGGGGAGCTCGCTGAGGCTTGACATGATTGGTGCGTA TGTTTG |
| AR prom | AR-Ble R | GAACGGCGCTGGTCAGCTTGGCCATTTAAGATGTTGAGTGAC |
| PSAD prom | pUC-PSAD F | TGACCATGATTACGAATTCGAGCTCGGTACATCCCACACACCTGCCCGTCTGCCTGACA |
| PSAD prom | pPSAD-Ble R | ACTGCTACTCACAACAAGCCCATGGCCAAGCTGACCAGCGCCGTT |
| METE prom | pUC-METE R | TGACCATGATTACGAATTCGAGCTCGGTACGTAGGTCAGGACCAGAGCCTACAAC |
| METE prom | pMETE-Ble R | CAGCTTGGCCATTTTAAGATGTTGAGTGACATGTCACTTAAATAATCGGCCTG |
| KG0-140 | BleGFP.Fw | TTGAAGACATAATGGCCAAGCTGACCAGC |
| KG0-140 | BleGFP.Rv | TTGAAGACATCGAACCCTTGTACAGCTCGTCCATG |

**Table S2.** List of plasmids generated by Gibson assembly.

| Plasmid | Terminator tested | Description | Figure |
|---|---|---|---|
| pMS3-0 | - | *pAR::Ble-GFP* | 3, 4, 5 |
| pMS3-8 | *RPL31* | *pAR::Ble-GFP::tRPL31* | 3 |
| pMS3-1 | *RPS29* | *pAR::Ble-GFP::tRPL29* | 3, 4 |
| pMS3-3 | *RPL11* | *pAR::Ble-GFP::tRPL11* | 3 |
| pMS3-14 | *RBCS2* | *pAR::Ble-GFP::tRBCS2* | 3 |
| pMS3-6 | *PSAD* | *pAR::Ble-GFP::tPSAD* | 3, 4, 5 |
| pMS3-11 | *THI4* | *pAR::Ble-GFP::tTHI4* | 3 |
| pMS3-10 | *METE* | *pAR::Ble-GFP::tMETE* | 3 |
| pMS3-12 | *CA1* | *pAR::Ble-GFP::tCA1* | 3, 4, 5 |
| pMS3-13 | *NIT1* | *pAR::Ble-GFP::tNIT1* | 3 |
| pMS3-N | *PSAD* | *pPSAD::Ble-GFP::tPSAD* | 5 |
| pMS3-K | *CA1* | *pPSAD::Ble-GFP::tPSAD* | 5 |
| pMS3-O | *PSAD* | *pMETE::Ble-GFP::tPSAD* | 5 |
| pMS3-L | *CA1* | *pMETE::Ble-GFP::tPSAD* | 5 |

**Table S3.** MoClo constructs employed and generated. The level 2 plasmids were used to generate data for Figure 6.

| Plasmid | Description | Function | Level | Source |
|---|---|---|---|---|
| pCM0-010 | *pPSAD (Pro + 5′UTR)* | Promoter + 5′UTR | 0 | [1] |
| pCM0-011 | *pAR (Pro + 5′UTR)* | Promoter + 5′UTR | 0 | [1] |
| pPM0-024 | *RBCS2i1 (5′UTR)* | RBCS2 intron 1 as 5′UTR enhancer | 0 | This study |
| pKG0-140 | *BleGFP (CDS)* | Resistance to zeocin + Reporter GFP | 0 | This study |
| pCM0-098 | *HA (C-ter tag, CDS)* | Immuno- and purification tag | 0 | [1] |
| pCM0-074 | *AphVIII (CDS)* | Resistance to paromomycin | 0 | [1] |
| pCM0-116 | *RSP29 (3′UTR + Ter)* | 3'UTR and Terminator | 0 | This study, [1] |
| pCM0-114 | *PSAD (3′UTR + Ter)* | 3'UTR and Terminator | 0 | [1] |
| pCM0-117 | *CA1 (3′UTR + Ter)* | 3'UTR and Terminator | 0 | This study, [1] |
| pFL_L1_004 | *pPSAD::RBCS2i1::BleGFPHA::tRPS29* | Expression cassette | 1 | This study |
| pFL_L1_005 | *pPSAD::RBCS2i1::BleGFPHA::tPSAD* | Expression cassette | 1 | This study |
| pFL_L1_006 | *pPSAD::RBCS2i1::BleGFPHA::tCA1* | Expression cassette | 1 | This study |
| pFL_L1_011 | *pAR::RBCS2i1::AphVIII::tPSAD* | Expression cassette | 1 | This study |
| pFL_L2_038 | *pPSAD::RBCS2i1::BleGFPHA::tRPS29::: pAR::RBCS2i1::AphVIII::tPSAD* | Expression cassette | 2 | This study |
| pFL_L2_044 | *pPSAD::RBCS2i1::BleGFPHA::tPSAD::: pAR::RBCS2i1::AphVIII::tPSAD* | Expression cassette | 2 | This study |
| pFL_L2_050 | *pPSAD::RBCS2i1::BleGFPHA::tCA1::: pAR::RBCS2i1::AphVIII::tPSAD* | Expression cassette | 2 | This study |

**Table S4.** Expression ranking of *C. reinhartii* genes used in this study. The expression rank over the diurnal cycle was determined by comparing the mean FPKM values in the dark, in the light and over the diurnal cycle. Data obtained from Strenkert *et al.* [2] (Dataset S2). The mean FPKM values (Mean), standard deviation (SD) and the expression rankings (Rank) are listed.

| Gene ID | Name | Expression in the dark[1] | | | Expression in the light[2] | | | Expression diurnal cycle[3] | | |
|---|---|---|---|---|---|---|---|---|---|---|
| | | Mean | SD | Rank | Mean | SD | Rank | Mean | SD | Rank |
| Cre12.g489153 | *L31* | 2023.91 | 176.15 | 40 | 2571.08 | 903.89 | 45 | 2425.76 | 803.35 | 42 |
| Cre08.g358556 | *RPS29* | 1312.28 | 125.52 | 82 | 1663.83 | 599.01 | 84 | 1597.48 | 563.75 | 79 |
| Cre01.g027000 | *RPL11* | 1578.22 | 131.49 | 61 | 1946.29 | 693.82 | 68 | 1889.14 | 651.09 | 63 |
| Cre02.g120150 | *RBCS2* | 5281.06 | 713.44 | 13 | 8078.13 | 2585.11 | 15 | 7054.93 | 2384.99 | 12 |
| Cre05.g238332 | *PSAD* | 974.70 | 349.65 | 116 | 2425.35 | 1012.79 | 51 | 1577.50 | 942.19 | 80 |
| Cre04.g214150 | *THI4* | 463.50 | 157.47 | 188 | 736.43 | 520.54 | 179 | 522.81 | 375.75 | 194 |
| Cre03.g180750 | *METE* | 39.82 | 90.98 | 812 | 396.42 | 176.57 | 232 | 173.57 | 214.41 | 345 |
| Cre04.g223100 | *CA1* | 42.56 | 44.69 | 771 | 417.34 | 343.95 | 225 | 217.22 | 268.27 | 300 |
| Cre09.g410950 | *NIT1* | 0.14 | 0.14 | 14971 | 0.13 | 0.04 | 15373 | 0.13 | 0.05 | 15537 |

[1] For expression in the dark, FPKM values from timepoints -11,-9,-7,-5,-3,-1h were used.
[2] For expression in the light, FPKM values from timepoints 1,3,5,7,9,11h were used.
[3] For expression over the diurnal cycle FPKM values from all timepoints were used.
